# Insights into transcriptional expression and putative functions of multiple polyhydroxyalkanoate synthase paralogs in *Haloferax mediterranei*

**DOI:** 10.1101/2025.07.03.663059

**Authors:** Chloé Vanden Haute, Brendan Schroyen, Ulrich Hennecke, Eveline Peeters

## Abstract

The halophilic archaeon *Haloferax mediterranei* is a promising candidate for polyhydroxyalkanoate production, offering several advantages due to its extremophilic physiology. While its primary polyhydroxyalkanoate synthase, a class III enzyme composed of PhaC_Hme_ and PhaE_Hme_ subunits, has been well characterized, the genome encodes three additional *phaC* paralogs (*phaC*_*1*_, *phaC*_*2*_ and *phaC*_*3*_), which were previously labeled as cryptic and remain poorly understood. In this study, we systematically investigated these paralogs by employing a targeted bioinformatics pipeline, revealing notable diversity in polyhydroxyalkanoate synthases among *Halobacteriales* and underscoring the distinctiveness of *H. mediterranei*. We further analyzed the native transcriptional expression profiles of all *phaC* paralogs under three physiologically relevant conditions: growth-limiting and growth-permissive conditions, as well as valeric acid supplementation to alter polyhydroxyalkanoate monomer composition. RT-qPCR analysis demonstrated that all three paralogs are transcriptionally active and differentially expressed, refuting earlier assumptions of their cryptic nature. Expression patterns were found not to correlate to polymer composition but to be dependent on growth phase, suggesting a potential physiological role for each paralog in native polyhydroxyalkanoate metabolism. These findings offer new insights into the functional complexity of polyhydroxyalkanoate biosynthesis in *H. mediterranei* and lay the groundwork for future metabolic engineering aimed at optimizing biopolymer production.

## Introduction

The search for sustainable alternatives to petrochemical polymers has directed growing interest towards bioplastics such as polyhydroxyalkanoates (PHAs). These biodegradable polyesters possess thermomechanical properties comparable to polypropylene and polyethylene (Laycock *et al*., 2013) and are produced by microbial biosynthesis. Beyond serving as carbon and energy storage (Dawes & Senior, 1973), PHAs can enhance host tolerance to environmental stress, such as ultraviolet irradiation, desiccation and osmotic stress (Tal & Okon, 1985), although the underlying mechanisms remain incompletely understood (Obruca *et al*., 2018).

Despite the potential, microbial PHA production remains several times more expensive than petrochemical counterparts (Tan *et al*. 2021). Extremophilic hosts have been proposed as a strategy to improve cost efficiency, offering advantages in fermentation and downstream processing (Kalia *et al*., 2025). Among them, the extremely halophilic archaeon *Haloferax mediterranei* has emerged as a promising candidate: its use as a production host can reduce costs by over 20% (Chen *et al*., 2022) through reduced sterility requirements and simplified cell lysis by hypo-osmotic shock in fresh water (Chen *et al*., 2006; Hermann-Kraus *et al*., 2013). As an archaeon, it also produces endotoxin-free PHAs, an advantage over bacterial hosts such as *Cupriavidus necator* in biomedical applications (Koller *et al*., 2013). Moreover, *H. mediterranei* grows on diverse low-cost carbon sources including dairy whey, glycerol from the biodiesel industry, vinasse and stillage from the bioethanol industry, chitin and crop residues (Koller *et al*., 2007a; Hermann-Krauss *et al*., 2013; Bhattacharyya *et al*., 2012; Bhattacharyya *et al*., 2014; Hou *et al*., 2013; Alsafadi *et al*., 2020).

A notable feature of *H. mediterranei* is its natural ability to synthesize poly(3-hydroxybutyrate-co-3-hydroxyvalerate) (PHBV) (Don *et al*., 2006). PHBV offers superior material properties compared to the more common poly(3-hydroxybutyrate) (PHB) homopolymer, being less crystalline, more flexible and having a larger processing window (Luzier, 1992). Unlike most production hosts, which require the addition of costly precursor molecules to synthesize PHBV, *H. mediterranei* synthesizes the copolymer directly from unrelated carbon sources (Han *et al*., 2013).

The key enzyme in the PHA synthesis pathway of *H. mediterranei* is a class III PHA synthase consisting of PhaE_Hme_ and PhaC_Hme_ (Lu *et al*., 2008). Genome sequencing has revealed three additional *phaC* paralogs: *phaC*_*1*_, *phaC*_*2*_ and *phaC*_*3*_ (Han *et al*., 2010). Under previously investigated PHA-accumulating conditions, none of the three paralogous *phaC* genes were transcribed and they were thus labeled as cryptic (Han *et al*., 2010), consistent with the observation that deletion of *phaC*_*Hme*_ completely abolished PHA synthesis (Lu *et al*., 2008). However, heterologous expression studies in *Haloarcula hispanica* indicated that PhaC_1_ and PhaC_3_, but not PhaC_2_, can form functional complexes with PhaE_Hme_, producing polymers with varying 3HV content (Han *et al*., 2010). Furthermore, deletion of a gene encoding an unusual phosphoenolpyruvate synthetase-like protein and located in a divergent operon with *phaC*_*1*_, induced the expression of *phaC*_*1*_ and *phaC*_*3*_, leading to the synthesis of PHA polymers with distinct monomeric compositions (Chen *et al*., 2020). These findings suggest that the paralogs could potentially be exploited for generating tailor-made copolymers in *H. mediterranei*.

Despite the presence of four *phaC* paralogs, of which minimally three have a functional capability, the extent to which cultivation conditions influence their transcription remains unclear, and a comparative structure–function analysis has not yet been performed. In this study, we investigate the role of PhaC_Hme_ and its paralogs in native PHA biosynthesis of *H. mediterranei*. By characterizing transcriptional expression profiles and the chemical composition of PHBV polymers in different growth conditions and phases, as well as modeling structural features of the corresponding enzymes, we aim to contribute to fundamental insights to support the rational design of strategies for production of customized PHAs.

## Materials and methods

### Bioinformatic analysis

Genomic neighbourhoods were analyzed using the accession numbers and genome assembly provided by NCBI. Protein structures were modeled using AlphaFold 3 (Abramson *et al*., 2024). Multiple sequence alignments were generated with COBALT (Papadopoulos *et al*., 2007).

To develop the bioinformatics pipeline, all *H. mediterranei* PhaC proteins (WP_004056138.1, WP_004060157.1, WP_004061055.1, WP_004060696.1) and the associated PhaE_Hme_ subunit (WP_004056139.1) were retrieved from the NCBI RefSeq database and used as reference queries. A dataset of PhaC-containing target species within the order *Halobacteriales* was compiled by extracting unique scientific names from an initial set of BLAST hits. Proteomes were downloaded using the NCBI Entrez API and concatenated to generate a single reference database representing *Halobacteriales*. A local BLAST database was constructed using the makeblastdb utility from BLAST+ (v2.16.0). Protein–protein BLAST (BLASTp) was run using each of the four PhaC reference sequences against the custom database, with the top four hits retained per query (-max_target_seqs 4) (Camacho *et al*., 2009). Output was formatted with extended fields (qacc sacc pident length mismatch gapopen qstart qend sstart send evalue bitscore stitle) to allow in-depth post-processing. A separate BLASTp search was conducted with PhaE_Hme_ (WP_004056139.1) as the query against the same database to assess genomic colocalization of PhaC and PhaE. BLAST results were parsed in Python (v 3.11) using pandas. Duplicate hits were removed based on identical alignment statistics across multiple fields and for each reference class (PhaC_Hme_, PhaC_1_, PhaC_2_, PhaC_3_), the highest scoring unique hit per organism was retained. The identity of the best matching reference for each hit was inferred by selecting the class with the highest percent identity. A hit was defined as weak when the percent identity of the best match was below 65%. To examine truncation patterns, the alignment length was compared to the full length of the reference sequence, expressed as a percentage of query coverage. To determine the genomic context of the genes corresponding to each PhaC and PhaE hit, genomic coordinates were retrieved using the NCBI Entrez efetch API. These coordinates were used to identify the closest PhaC-encoding gene for each hit encoding PhaE within the same genome, based on the smallest absolute distance between the end coordinate of *phaE* and the start of the corresponding *phaC* gene. This information was used to infer co-localized gene pairs and assign each PhaE to its most likely functional PhaC partner.

Strong hits (percent identity ≥ 65%) were classified as either PhaC_Hme_-, PhaC_1_-, PhaC_2_-or PhaC_3_-like based on identity scores and visualized in Python (v 3.11) using matplotlib. To assess patterns of paralog occurrence, summary statistics were calculated from the final dataset. Species with one or more PhaC paralogs were tallied, and the most prevalent number of paralogs was identified. The distribution of best-matching classes was examined. Additionally, the number of species for which a PhaC-PhaE pairing could be confidently established based on genomic colocalization was quantified. Each strong PhaC hit was represented as a colored circle according to its best matching class, plotted per organism and vertically stacked.

### Strains, media and cultivation conditions

*H. mediterranei* DSM1411 was cultivated at 42°C and 125 rpm in an orbital shaker in different liquid defined media derived from the Hv-min minimal medium (Dyall-Smith *et al*., 2009). Briefly, this medium contained 15.9 mM Tris-HCl (pH 7.5), 2.6 M NaCl, 93.8 mM MgCl_2_, 90.3 mM MgSO_4_, 60 mM KCl, 5.3 mM NH_4_Cl, 6.4 mM CaCl_2_, 0.6 mM K_2_HPO_4_, 0.4 mM KH_2_PO_4_, 1.9 µM MnCl_2_, 1.6 µM ZnSO_4_, 8.8 µM FeSO_4_, 0.2 µM CuSO_4_, 3.2 µM thiamine and 0.4 µM biotin. This medium was supplemented either with 32.0 mM sodium DL-lactate, 3.7 mM glycerol and 15.0 mM sodium succinate dibasic hexahydrate (labelled “Hv-min”) or only with 74.2 mM glycerol (labelled “glycerol medium”) as carbon source. Growth was monitored by measuring optical density at 600 nm (OD_600_). When mentioned, the Hv-min medium was supplemented with 0.5 g L^-1^ valeric acid in exponential phase (OD_600_ between 0.4 and 0.6).

### Nucleic acid extractions and cDNA preparation

For genomic DNA (gDNA) extraction, *H. mediterranei* cells were harvested by centrifugation at 16,000 x *g* during 2 minutes followed by resuspension in Elution Buffer (“EB” 10 mM Tris-HCl pH 8.5) and addition of an equal volume of phenol/chloroform/isoamyl alcohol solution (25:24:1). This solution was vortexed, centrifuged at 16,000 x *g* during 5 minutes, the top aqueous phase was recovered and this procedure was repeated. Next, an equal volume of chloroform/isoamyl alcohol solution (24:1) was added and the top aqueous phase was removed in the same manner. This phase was combined with 0.75 M NH_4_OAc and 20 µg glycogen, followed by ethanol precipitation after which the DNA-containing pellet was resuspended in EB.

RNA was extracted using a RNeasy Mini Kit (Qiagen) with two consecutive DNase treatments according to manufacturer’s instructions. The starting culture volumes for extractions at different growth phases were normalized to the OD_600_, in order to start from similar amounts of cells. RNA was converted to cDNA using a GoScript^TM^ Reverse Transcriptase kit (Promega) following manufacturer’s instructions.

### Gene expression analysis by quantitative RT-PCR

Specific primers for quantitative RT-PCR (RT-qPCR) were designed employing Primer3 (Untergasser *et al*., 2012) (**Supplementary Table S1**). For each primer pair, amplification efficiency and primer specificity was verified employing a ten-fold dilution series of gDNA as a template. RT-qPCR reactions were performed with GoTaq qPCR Master Mix (Promega) and 10 µM of each primer in a CFX96 Real-Time System (Bio-Rad). Each condition was analyzed using three biological replicates in two technical replications. The following thermal cycling program was performed: 3 minutes at 95°C followed by 40 cycles of 10 seconds at 95°C and 30 seconds at 55°C.

The Pfaffl method was used to calculate relative gene expression ratios (Pfaffl, 2001). As a housekeeping gene, either *tbp* (E6P09_03900) or *ffs* (E6P09_08165) was used, or the geometric means of both (Vandesompele *et al*., 2002, **Supplementary Note S1**). Statistical analysis was performed by averaging technical replicates and comparing the resulting groups of three biological replicates by means of an unpaired Student’s t-test in the Prism 10 software package (GraphPad).

Quantitative transcript levels were estimated by relating the measured Ct values of RT-qPCR assays to the standard curves recorded for each gene (**Supplementary Figure S1** and **Supplementary Table S2**). The DNA quantities were then used as a proxy for transcript levels, allowing the comparison of different paralogs at different timepoints.

### PHA extraction

*H. mediterranei* cells were harvested in stationary phase (OD_600_ around 2.0 for Hv-min, 2.7 for Hv-min with valeric acid and around 0.5 for glycerol medium), centrifugated at 12,100 x *g* during 15 minutes, washed with HSPB buffer (2 M NaCl, 71.7 mM K_2_HPO_4_ and 28.3 mM KH_2_PO_4_) and snap-frozen in liquid nitrogen. When frozen, the cells were lyophilized for 16 hours in a benchtop lyophilizer (Labconco). The dried pellets were weighed and placed in a Whatmann Extraction Thimble, which was suspended in 30 mL ethanol and incubated at room temperature while shaking for 16 hours. The thimble was transferred to 30 mL chloroform and incubated. Lastly, chloroform was allowed to evaporate overnight, after which the resulting PHA was weighed.

### Nuclear Magnetic Resonance analysis

The monomeric composition of the extracted PHA polymers was determined by ^1^H-NMR spectroscopy. Polymers were dissolved in deuterated chloroform (CDCl_3_) and ^1^H-NMR spectra were measured at 400 MHz on a Bruker Avance Neo spectrometer (Bruker, Germany) using a standard pulse sequence at ambient temperature. The chemical shifts are reported in delta (δ) units in parts per million (ppm) relative to trimethylsilane. Spectra were referenced relative to the residual solvent signal. The 3-hydroxyvalerate content of the polymers was calculated from the ratio of the area under the methine proton peaks of the 3-hydroxybutyrate and 3-hydroxyvalerate subunits, respectively.

## Results

### Comparative analysis of PhaC paralogs in H. mediterranei

All four *phaC* paralogous genes are located at distinct genomic loci in *H. mediterranei*, including the chromosome and extrachromosomal elements (**Figure 1A**). While *phaC*_*Hme*_, which is considered to encode the main PhaC enzyme in *H. mediterranei*, is located on the pHM322 megaplasmid in an operon with the gene encoding its auxiliary protein PhaE_Hme_ (Lu *et al*., 2008) adjacent to another operon with other PHA-related genes, the other paralogs are encoded on genomic locations without any PHA-related genes in their neighbourhood. Whereas *phaC*_*1*_ is located on the chromosome, the *phaC*_*2*_ and *phaC*_*3*_ genes are encoded on the megaplasmid pHME505 (**Figure 1A**).

**Figure 1.**
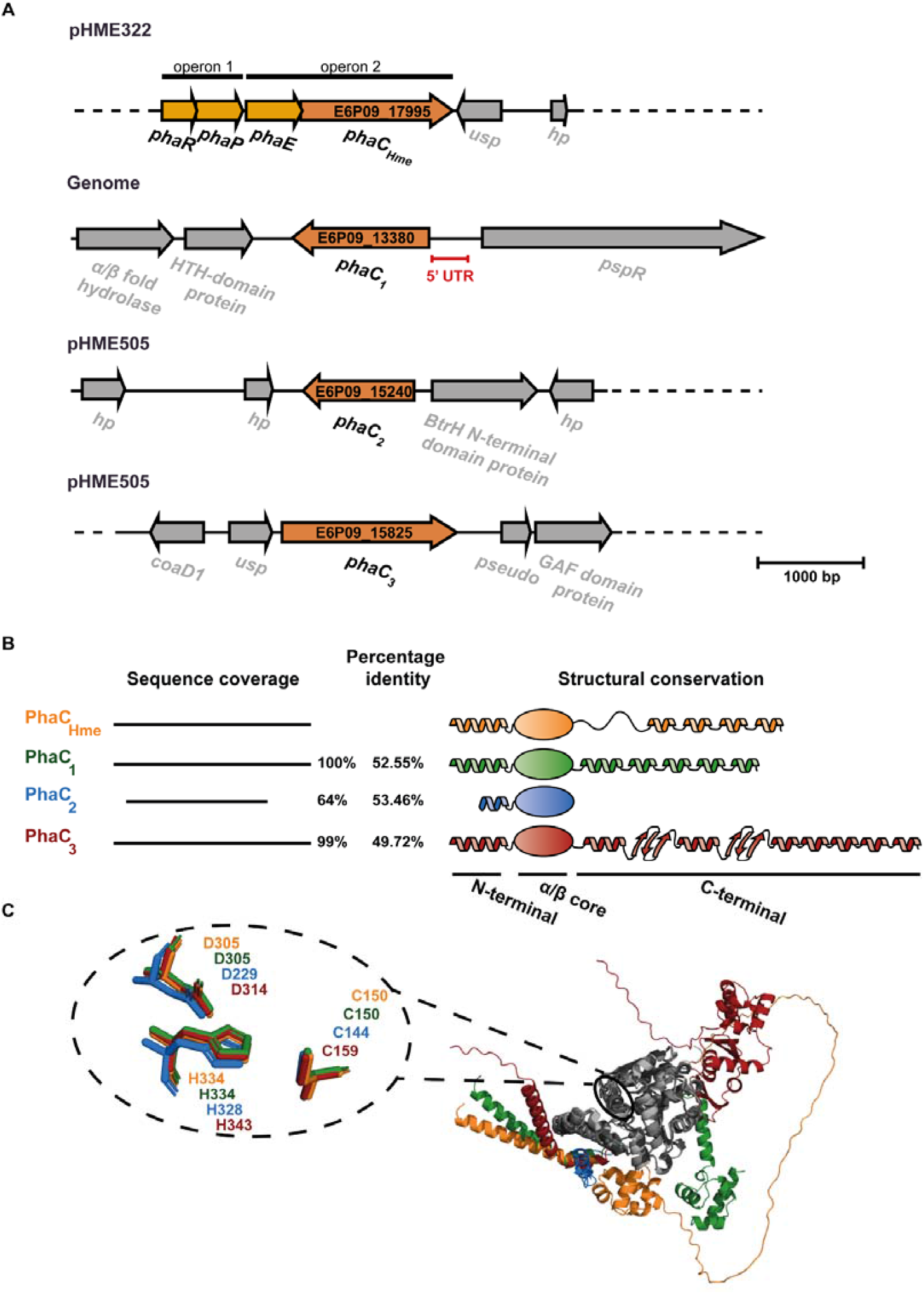
Genome organization of *phaC* paralogs in *H. mediterranei* and sequence and structural similarities of the resulting proteins. **A**. Schematic overview of the genome organization of the four *phaC* paralogs in *H. mediterranei*. Usp: universal stress protein, hp: hypothetical protein, HTH: helix-turn-helix, *pspR*: PPS-like regulator, BtrH: Butirosin biosynthesis protein H, *coaD1*: phosphopantetheine adenylyltransferase. **B**. Sequence coverage with respect to the PhaC_Hme_ sequence, with indication of sequence identities of PhaC_1_, PhaC_2_ and PhaC_3_ with respect to PhaC_Hme_. On the right, a schematic representation is shown of the domain organization of the PhaC enzymes, with emphasis on their secondary structure elements. **C**. Structural superposition of the catalytic cores of the four enzymes, with a zoomed view of the three catalytically indispensable residues and their respective positions within the PhaC structures.

Based on a multiple sequence alignment (**Supplementary Figure S2**), it can be observed that PhaC_1_ and PhaC_3_ cover nearly full sequence length of the primary enzyme PhaC_Hme_, while PhaC_2_ only covers 62% of the sequence, missing segments at the C- and N-termini (**Figure 1B**). All paralogs exhibit between 44% and 60% sequence identity (**Supplementary Figure S3**). Structural modeling confirmed with high confidence that the conserved central segment of the proteins corresponds to the α/β core, which is a signature for the α/β hydrolase superfamily (**Figure 1C; Supplementary Figure S4**). Within this core, the indispensable catalytic triad, consisting of a histidine, cysteine and aspartate residue, is positionally conserved (**Figure 1C**). In contrast to the core of the enzyme, structural features of the termini are predicted with a lower confidence for all paralogs, in agreement with a lower conservation. While all paralogs are predicted to form an α helix at the N-terminus, albeit being much smaller for PhaC_2_, its relative position varies significantly (**Figure 1B**). The N-terminal PhaC domain is hypothesized to be involved in different types of interactions including protein-granule interactions and the formation of dimers or other protein-protein interactions (Neoh *et al*., 2022). A large variability is observed for the C-terminal region, which is called the CAP domain and functions to close the active site and to block the substrate entry path (Chek *et al*., 2017). Whereas PhaC_2_ lacks a C-terminal domain, PhaC_3_ is predicted to form an extended C-terminal domain with two additional β sheets. The absence of the CAP domain in PhaC_2_ might render the enzyme non-functional as previously shown (Han *et al*., 2010).

### Phylogenetic distribution of PhaC in Halobacteriales

To determine whether the occurrence of multiple PhaC paralogs is a conserved trait in PHA-producing *Halobacteriales*, we screened genomes predicted to contain at least one identified *phaC* homolog that were thus considered putative PHA producers (**Figure 2**). Of 225 genomes, only a limited number harbor more than one *phaC* gene: 32 species were predicted to encode two paralogs, 4 species to encode three, while *H. mediterranei* is the only species with four paralogs.

**Figure 2.**
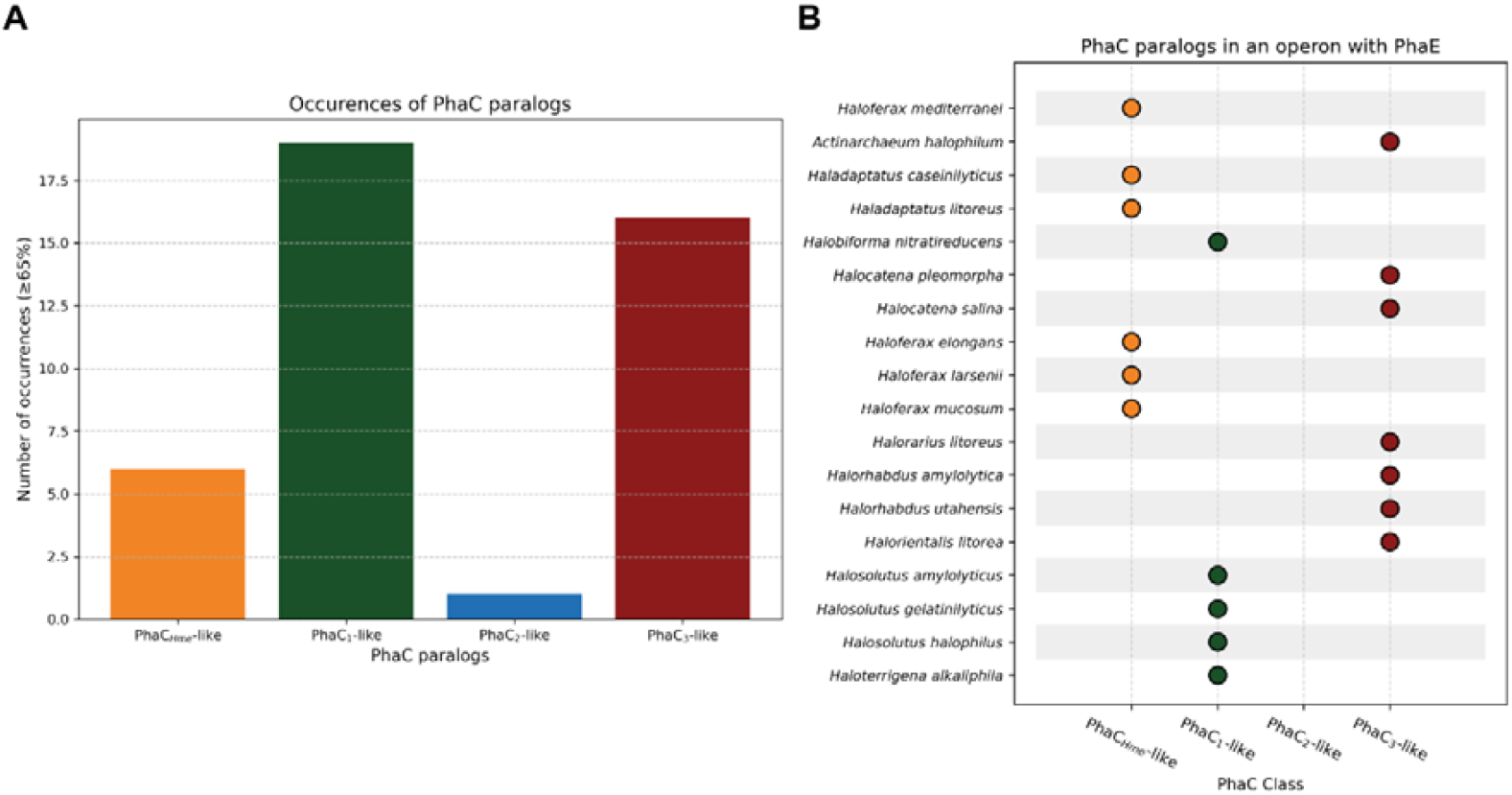
Graphical representation of the occurrence of PhaC paralogs in the genomes of PHA-producing *Halobacteriales*. PhaC homologs were classified into four categories—PhaC_Hme_-like, PhaC_1_-like, PhaC_2_-like or PhaC_3_-like —based on sequence identity. **A**. Bar chart representing the distribution of PhaC-like paralogs in *Halobacteriales*. **B**. Overview of PhaC-like paralogs encoded in the vicinity of a PhaE subunit, suggesting the presence of a *phaEC* operon.

Upon comparing the detected PhaC sequences with each of the *H. mediterranei* paralogs, a scattered phylogenetic occurrence was observed, where the majority showed the highest sequence identity with either PhaC_1_ or PhaC_3_ of *H. mediterranei* (**Figure 2A**). In contrast, C-terminal truncated PhaC_2_-like homologs were not detected in any other species, suggesting that this paralog may be a unique evolutionary feature in *H. mediterranei*. In addition to *phaC* diversity, patterns in *phaE* co-localization were examined (**Figure 2B**). It was observed that the gene encoding the PhaE subunit frequently co-occurs in the genome with various *phaC* paralogs, not exclusively with one particular class. Despite the presence of multiple *phaC* genes in some genomes, most species contained only a single copy of *phaE*. Only in a few exceptional cases, for example in *Natronomonas aquatica*, more than one *phaE* gene was identified.

### Transcriptional expression of phaC paralogs in different medium conditions

To investigate whether the four *phaC* paralogs are transcriptionally expressed and how their expression changes with different carbon sources, we selected two defined media: i) Hv-min medium, containing a mixture of glycerol, sodium lactate and sodium succinate and ii) glycerol medium, containing only glycerol (**Figure 3**). First, these media were compared for their ability to support growth and PHA production in *H. mediterranei*. While the glycerol medium supported growth to a maximal OD_600_ of 0.533 ± 0.062, which is relatively low for this species, the Hv-min medium reached a much higher carrying capacity with a maximal OD_600_ of 2.220 ± 0.003 (**Figure 3A**). It can thus be concluded that glycerol medium represents a growth-limiting condition and Hv-min medium a growth-permissive condition. In terms of PHA production, both media performed similarly. NMR analysis (**Supplementary Figure S5** and **S6**) demonstrated that the PHBV composition was comparable in both growth conditions with 10.4 ± 1.3 % 3-hydroxyvalerate (3HV) and 9.2 ± 0.6 %3HV in the glycerol and Hv-min medium, respectively (**Figure 3B**). The PHA content was relatively low, namely 5.2 ± 1.8 % in glycerol medium and 6.5 ± 1.5% in Hv-min medium (**Figure 3C**). Despite large differences in biomass yield, both media supported similar PHA production, suggesting that PHA synthesis in *H. mediterranei* is largely independent of growth rate. These two media therefore provide suitable conditions to investigate *phaC* expression and its regulation in a comparative manner, as PHA synthesis is relevant in both conditions, yet the cells are in markedly different metabolic and physiological states.

**Figure 3.**
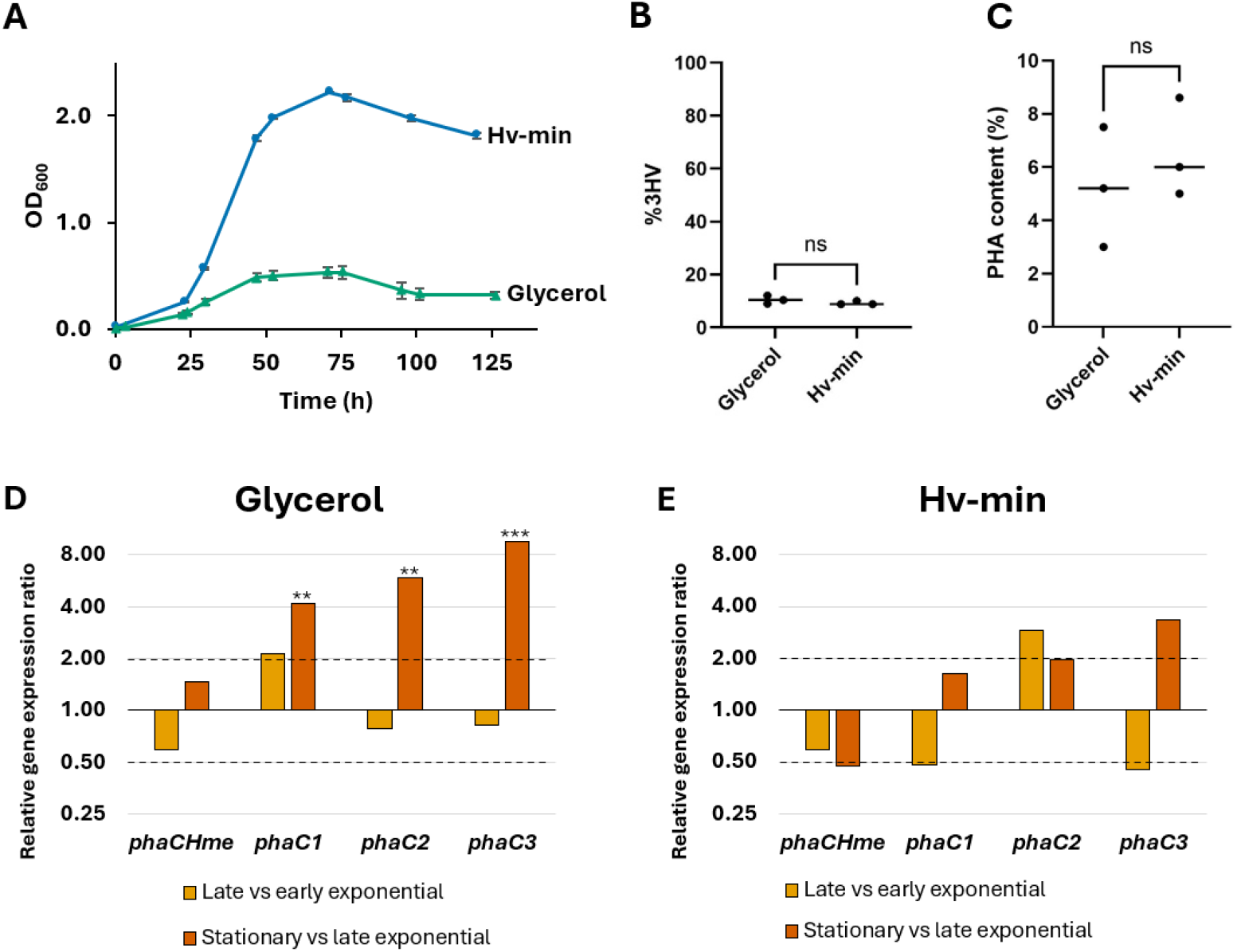
Growth, PHA synthesis and relative gene expression of *phaC* paralogs in growth-limiting and - permissive conditions. **A**. Growth curves of *H. mediterranei* in glycerol medium and Hv-min defined media, measured as optical density at a wavelength of 600 nm (OD_600_) over time. Values represent the average of three biological replicates and error bars indicate standard deviations. **B**. 3-Hydroxyvalerate (%3HV) content in the PHA polymers produced in glycerol and Hv-min media, as measured by NMR. Triplicate values were compared using a *t*-test; ns = not significant. **C**. PHA content in glycerol and Hv-min media. Triplicate values were compared using a *t*-test; ns = not significant. **D**. Relative transcriptional expression of *phaC* paralogs in growth-limiting conditions, as determined by RT-qPCR. Ct values were compared using a statistical *t*-test. ** indicates *p* = 0.01 - 0.001, *** indicates *p* < 0.001. Dotted lines indicate thresholds for 2.0 (upregulation) and 0.5 (downregulation). **E**. Relative transcriptional expression of *phaC* paralogs in growth-permissive conditions, as determined by RT-qPCR.

In both the growth-limiting (glycerol medium) and growth-permissive (Hv-min medium) condition, transcriptional expression levels of *phaC*_*Hme*_, *phaC*_*1*_, *phaC*_*2*_ and *phaC*_*3*_ were monitored employing RT-qPCR (**Figure 3D** and **E**). In contrast to previous claims that *phaC*_*1*_, *phaC*_*2*_ and *phaC*_*3*_ are cryptic genes (Han *et al*., 2010; Chen *et al*., 2020), we observed transcriptional activity for all *phaC* paralogs in these conditions. Moreover, in the growth-limiting condition, *phaC*_*1*_, *phaC*_*2*_ and *phaC*_*3*_ displayed a significant transcriptional upregulation when comparing stationary to late exponential growth phase (**Figure 3D**). A slight upregulation was already apparent for *phaC*_*1*_ upon comparing late to early exponential growth phase. In contrast, the expression level of *phaC*_*Hme*_ remained relatively constant over all investigated timepoints. Although less pronounced, a similar trend of upregulation towards the stationary growth phase was observed in the growth-permissive condition (**Figure 3E**), which was most pronounced for *phaC*_*2*_ and *phaC*_3_. Simultaneously, *phaC*_*Hme*_ displayed a slight downregulation in later growth stages. Altogether, these observations suggest that in later growth phases, *phaC*_*1*_, *phaC*_*2*_ and *phaC*_*3*_ have a relatively more important role in PHA synthesis than the traditionally recognized key gene *phaC*_*Hme*_.

### Effects of valeric acid supplementation on transcriptional expression of phaC paralogs

Although *H. mediterranei* is inherently capable of producing PHBV (**Figure 3B**), previous studies have shown that the 3HV content of the copolymer can be increased by supplementing the medium with a precursor such as valeric acid (Ferre-Guell & Winterburn, 2018). Here, we supplemented exponentially growing cells under growth-permissive conditions (Hv-min medium) with 0.5 g L^-1^ valeric acid and monitored its impact on growth and PHA synthesis (**Figure 4**). The addition of valeric acid stimulated growth, prolonging the exponential phase and resulting in a higher OD_600_ (2.990 ± 0.090 as compared to 2.220 ± 0.003) (**Figure 4A**). The addition of valeric acid significantly increased the 3HV content in the polymer of *H. mediterranei* from 9.2 ± 0.6% to 53.4 ± 3.7% (**Figure 4B**; **Supplementary Figure S7**). Moreover, precursor feeding boosted the PHA content from 6.5 ± 1.5% to 10.5 ± 1.2% (**Figure 4D**).

**Figure 4.**
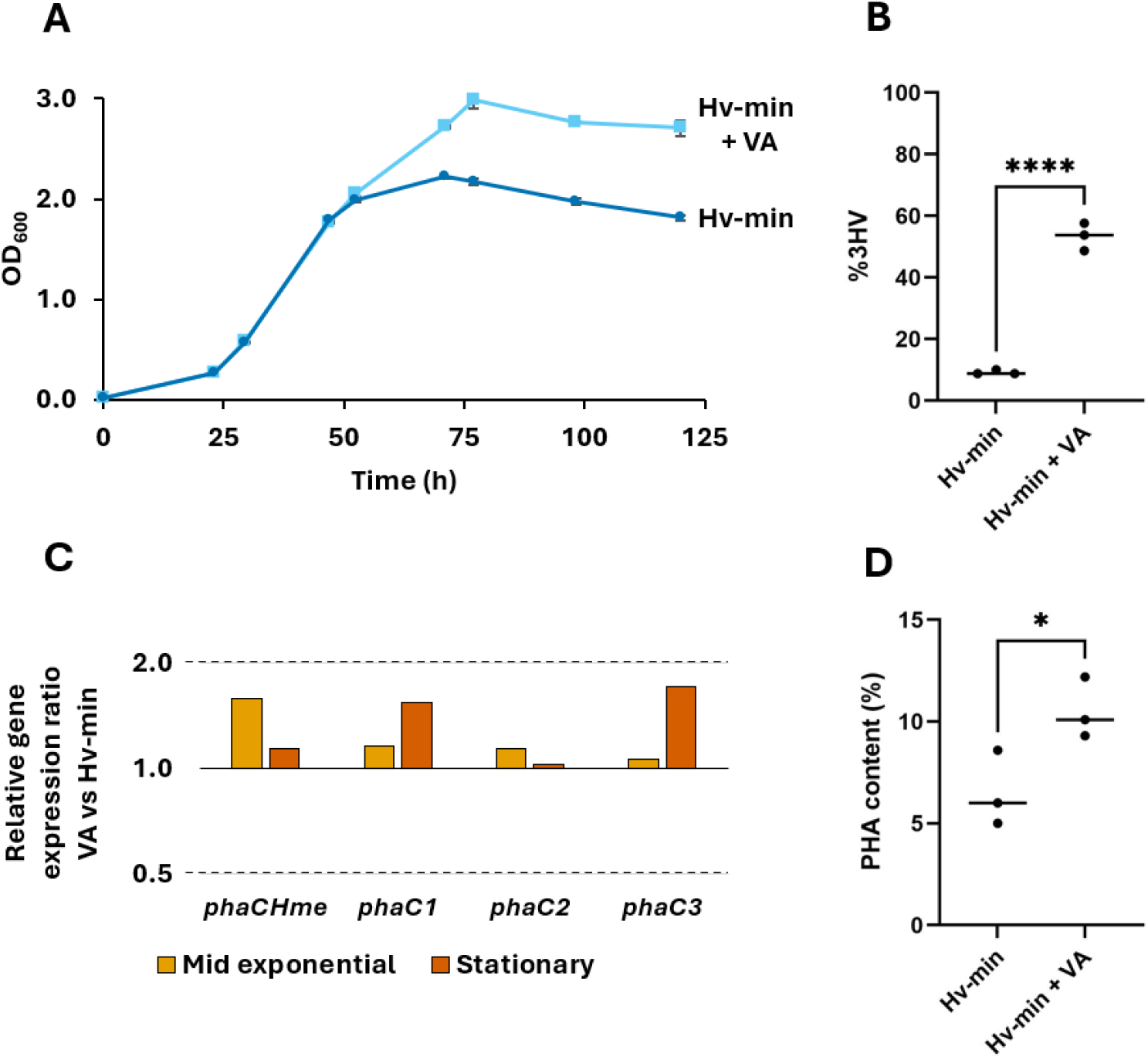
Growth, PHA synthesis and relative gene expression of *phaC* paralogs upon addition of valeric acid. **A**. Growth curves of *H. mediterranei* in Hv-min medium with and without valeric acid supplementation, measured as optical density at 600 nm (OD_600_) over time. Values represent the average of three biological replicates; error bars indicate standard deviations. **B**. 3-Hydroxyvalerate (%3HV) content in the PHA polymers produced in Hv-min medium with (Hv-min + VA) and without valeric acid (Hv-min), as measured by NMR. Triplicate values were compared using a *t*-test; **** = *p* < 0.0001. **C**. Relative transcriptional expression of *phaC* paralogs in Hv-min medium comparing addition of valeric acid to the absence of it, as determined by RT-qPCR. Dotted lines indicate thresholds for 2.0 (upregulation) and 0.5 (downregulation). **D**. PHA content in Hv-min medium with and without valeric acid suplementation. Triplicate values were compared using a *t*-test; * indicates *p* = 0.05 - 0.01.

Given that PhaC_1_ and PhaC_3_ were previously shown to synthesize PHAs with distinct monomeric compositions upon heterologous expression (Han *et al*., 2010), we investigated the transcriptional gene expression levels of all *phaC* paralogs in Hv-min medium with and without valeric acid supplementation (**Figure 4C**). No significant differential expression was observed in either the mid-exponential or stationary growth phase. This indicates that transcriptional regulation of the paralogs is not responsive to the presence of a 3HV precursor and that it is independent of the copolymer’s 3HV content.

The growth phase dependence of gene expression of the *phaC* paralogs was also investigated in precursor-supplemented medium by comparing the stationary and mid-exponential growth phase. The three paralogs *phaC1, phaC2* and *phaC3* again showed significant upregulation in the stationary phase (relative expression ratios of 3.69, 2.55 and 5.68 for *phaC1, phaC2* and *phaC3*, respectively) while the main subunit *phaC*_*Hme*_ showed a downregulation (relative gene expression ratio of 0.33) (**Figure 5**). These results further corroborate an increased importance of the *phaC1, phaC2* and *phaC3* paralogs in later growth stages.

**Figure 5.**
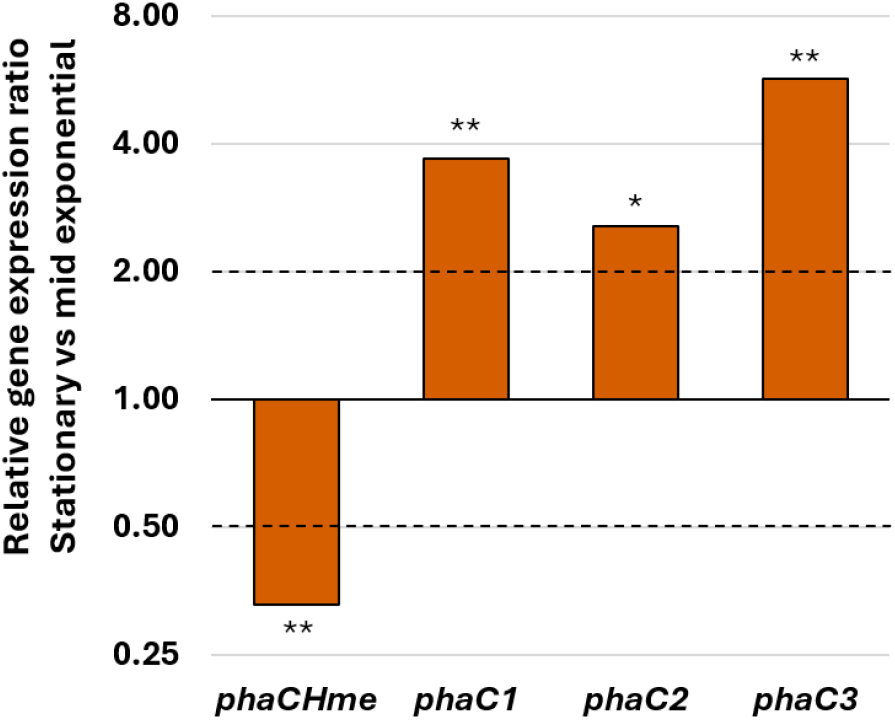
Relative gene expression of *phaC* paralogs in stationary phase relative to mid-exponential phase upon addition of valeric acid. Relative transcriptional expression was determined by RT-qPCR. Ct values were compared using a statistical *t*-test. * indicates *p* = 0.05 - 0.01, ** indicates *p* = 0.01 - 0.001. Dotted lines indicate thresholds for 2.0 (upregulation) and 0.5 (downregulation).

### Differences in relative transcription levels of the phaC paralogs

Given that all paralogs are actively transcribed under the investigated conditions, we next set out to analyze potential differences in relative transcription levels of the *phaC*_*1*_, *phaC*_*2*_ and *phaC*_*3*_ paralogs compared to the main *phaC*_*Hme*_ paralog in different growth phase stages (**Figure 6**). Using standard curves generated with gDNA as a template, measured Ct values were converted into input DNA quantities, which were then used as an approximation of transcript levels. Throughout the growth process, *phaC*_*Hme*_ was transcribed at significantly higher levels than the three other paralogs (*phaC1, phaC2* and *phaC3*) across all investigated media (**Figure 6**). However, the difference in transcription levels decreased as growth progressed, with the gap between the main subunit and the other paralogs becoming less pronounced in the late exponential and stationary phases.

**Figure 6.**
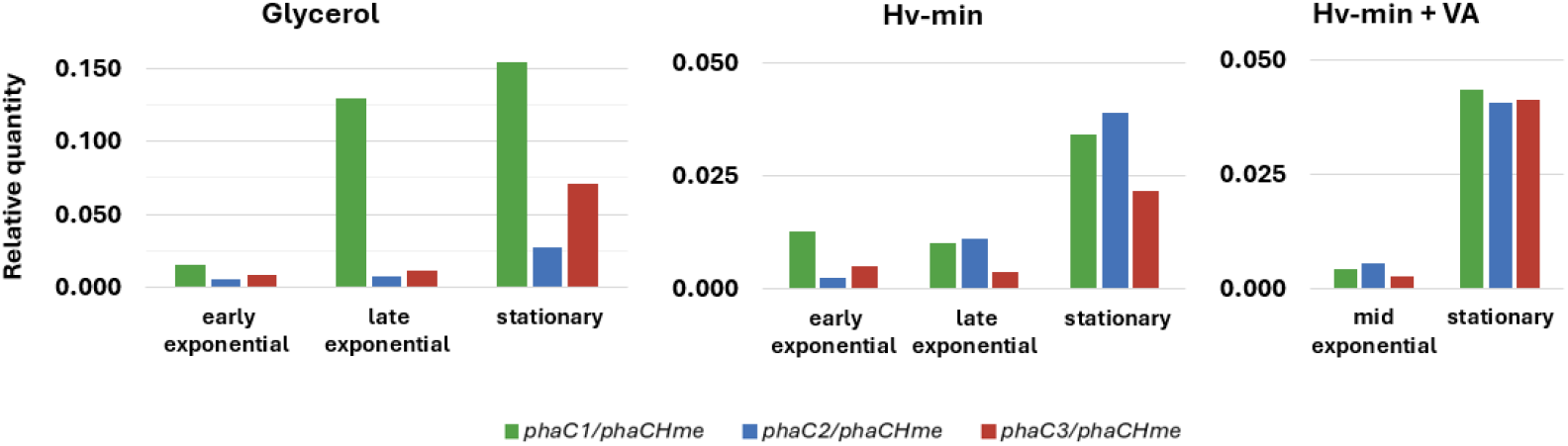
Relative quantity of transcript levels of the main *phaC* subunit (*phaCHme)* compared to each paralog in all investigated conditions. Values for each gene are derived by relating measured Ct values to their respective standard curves.

## Discussion

Our work has shown that *H. mediterranei* is the only species within the class *Halobacteriales* that encodes four distinct PhaC paralogs. Structural conservation across the catalytic cores of all four enzymes, combined with variations in the terminal regions, indicates a shared enzymatic ancestry with potentially divergent functional capacities. The consistent presence of a conserved N-terminal α-helix in all paralogs further suggests that interaction with PhaE remains possible. A screening of 225 *Halobacteriales* genome sequences revealed that approximately 17% are also predicted to encode multiple paralogs, though only two or three and predominantly PhaC_1_-like or PhaC_3_-like homologs. No other species was found to encode a homolog of PhaC_2_, which lacks the C-terminal CAP domain and was previously shown to be functionally inactive (Han *et al*., 2010). This suggests that PhaC_2_ is unique to *H. mediterranei*.

Whereas a previous study reported that only *phaC*_*Hme*_ was actively transcribed and that the three other paralogs represented cryptic genes (Han *et al*., 2010), our results demonstrate that all *phaC* paralogs are expressed under different PHA-producing growth conditions. In line with previous findings (Han *et al*., 2010; Yu *et al*., 2008), our results confirm the role of PhaC_Hme_ as the primary PhaC subunit in *H. mediterranei*, as it consistently exhibited the highest transcription levels. This conclusion is further supported by its genomic context, with *phaC*_*Hme*_ being co-located with *phaE*_*Hme*_ in a single operon, both being under the regulation of transcription factor PhaR (Lu *et al*., 2008). When looking beyond *Halobacteriales*, the occurrence of multiple PhaC paralogs is not unusual. In *Rhodospirillum rubrum*, several paralogs (PhaC_1_ & PhaC_3_) are functionally redundant and contribute little to PHA metabolism under standard growth conditions (Jin *et al*., 2012). In *Bradyrhizobium japonicum*, five PhaC paralogs are encoded in the genome of which only two (PhaC1 and PhaC2) are significantly expressed (Quelas *et al*., 2013). Although both are expressed at similar levels in wild-type cells, no PHA accumulation was observed in a ΔPhaC1 strain, whereas PHA production was increased in a ΔPhaC2 strain, indicating a complex interplay between the two enzymes. In the same study, it was hypothesized that the paralogs form a heterodimer and mutually balance out each other’s activity (Quelas *et al*., 2013).

As previous reports suggested that multiple PhaC paralogs could be involved in the synthesis of polymers with distinct monomeric compositions (Han *et al*., 2010; Chen *et al*., 2020), we assessed the effects of precursor addition on the transcriptional expression of the *phaC* genes in *H. mediterranei*. PHBV produced by this species typically contains about 10% 3HV, independent of the carbon source (Chen *et al*., 2006; Koller *et al*., 2007a; Bhattacharyya *et al*., 2012; Bhattacharyya *et al*., 2014), consistent with our findings on glycerol and Hv-min media. The potential of valerate as a 3HV precursor in *H. mediterranei* had already been investigated previously (Koller *et al*., 2007b; Han *et al*., 2015); however, our approach resulted in considerably higher 3HV incorporation at the tested precursor concentration compared to earlier reports (21.8% for Koller *et al*. at 8 mM valerate; 20.8% for Han *et al*. at 6.5 mM valerate; 53.4% in our study at 4.9 mM valerate). A possible explanation for this discrepancy is our use of valeric acid as the source of valerate, which might cause pH stress, thereby altering PHA production characteristics. Despite the large difference in HV content, we observed that the regulation of all *phaC* paralogs appears to be independent of PHA content or composition.

Although the relative expression levels of *phaC*_*1*_, *phaC*_*2*_ and *phaC*_*3*_ were markedly lower than that of the canonical *phaC*_*Hme*_, they were nonetheless regulated, indicating potential physiological relevance. This regulation was growth-phase-dependent, with *phaC*_*1*_ and *phaC*_*3*_ being significantly upregulated during the stationary phase, while expression of the main subunit remained constant or showed a slight downregulation, thereby gradually lowering the difference between the expression level of *phaC*_*Hme*_ and those of *phaC*_*1*_, *phaC*_*2*_ and *phaC*_*3*_. It is possible that transcripts of *phaC*_*1*_, *phaC*_*2*_ and *phaC*_*3*_ went undetected in the previous study due to transcription being assessed exclusively during the exponential growth phase (Han *et al*., 2010). However, the detection of transcription across all paralogs, with consistently higher levels of *phaC*_*Hme*_ compared to *phaC*_*1*_, *phaC*_*2*_ and *phaC*_*3*_ is consistent with values previously reported in an RNA-sequencing study, also in exponentional growth phase (Chen *et al*., 2019; Chen *et al*., 2020).

Interestingly, in other PHA-producing *Halobacteriales*, different paralog classes - except for true PhaC_2_-like variants - can be found within *phaEC*-like operons, corroborating the hypothesis that the functional relevance of the paralogs in *H. mediterranei* is influenced by regulated expression. The tight regulation of the non-canonical *phaC* paralogs suggests the existence of a broader regulatory network involving multiple *trans*-acting factors. This includes the phosphoenolpyruvate synthase-like protein PspR, whose gene is encoded in a divergent operon with *phaC*_*1*_ (**Figure 1A**). Deletion of *pspR* has been shown to correlate with higher transcriptional expression of *phaC*_*1*_ and *phaC*_*3*_, although the underlying mechanism are not entirely clear (Chen *et al*., 2020; Chen *et al*., 2024). Additional regulators might include PhaR and Fnr-like proteins (Mitra *et al*., 2021; Chen *et al*., 2024), and it can be hypothesized that the expression of the paralogs is also subject to post-transcriptional regulation. In this context, it is noteworthy that the *phaC*_*1*_ paralog possesses an exceptionally long 5’-UTR region (∼350 nt) (**Supplementary Figure S8**) (Martinez Pastor *et al*., 2025). Future research could focus on unraveling the underlying regulatory network, with the hypothesis that the non-canonical *phaC* paralogs may be upregulated under diverse stress conditions, not only stationary-phase growth, and could contribute to modulating the properties of the synthesized biopolymer.

## Funding

This work was supported by the Vrije Universiteit Brussel (Strategic Research Program SRP91). C.V.H. and B.S. were funded by PhD fellowships of the Research Foundation Flanders (FWO-Vlaanderen) with respective grant numbers [1SH8V24N] and [1S21424N]. U.H. is grateful for the financial support by the VUB/FWO grant OZR 3584 (NMR spectrometer).

## Author contributions

C.V.H.: Conceptualization, Investigation, Formal analysis, Writing – original draft. B.S.: Conceptualization, Investigation, Formal analysis, Writing – original draft. U.H.: Resources, Writing – review & editing. E.P.: Resources, Supervision, Writing – review & editing.

## Notes

### Competing Interest Statement

The authors have declared no competing interest.

### Summary of Updates

Revised to streamline text and adhere to the journal's policy

