## Supplementary for "Insights into transcriptional expression and putative functions of multiple polyhydroxyalkanoate synthase paralogs in *Haloferax mediterranei*"

### SUPPLEMENTARY MATERIALS - Insights into transcriptional expression and putative functions of multiple *phaC* paralogs in *Haloferax mediterranei*

**<u>Supplementary Note S1:</u>** Relative gene expression ratios (R, Equation 2) were calculated according to the Pfaffl method. Inputs to the Pfaffl equation are modified primer efficiencies (E, Equation 1) and Ct values of both the target genes and a housekeeping gene under a sample and a control condition. For the analysis over different growth stages in the glycerol and the Hv-min+VA media, *tbp* was used as the stable housekeeping gene whereas *ffs* was used in Hv-min medium. For comparison of the Hv-min and Hv-min+VA conditions, the geometric mean of both *tbp* and *ffs* was used according to Equation 3. All input Ct values were the average of two technical replicates and three biological replicates.

$$E = 1 + \left[\frac{amplification\ efficiency\ (\%)}{100}\right] \tag{1}$$

$$R = \frac{{E_{target}}^{\Delta Ct\ (control-sample)}}{{E_{HK}}^{\Delta Ct\ (control-sample)}} \tag{2}$$

$$R = \frac{{E_{target}}^{\Delta Ct\ (control-sample)}}{GeoMean\left[{E_{HK}}^{\Delta Ct\ (control-sample)}\right]} = \frac{{E_{target}}^{\Delta Ct\ (control-sample)}}{\prod_{i=1}^{n}\left[{E_{HK}}^{\Delta Ct\ (control-sample)}\right]_i^{\frac{1}{n}}} \tag{3}$$

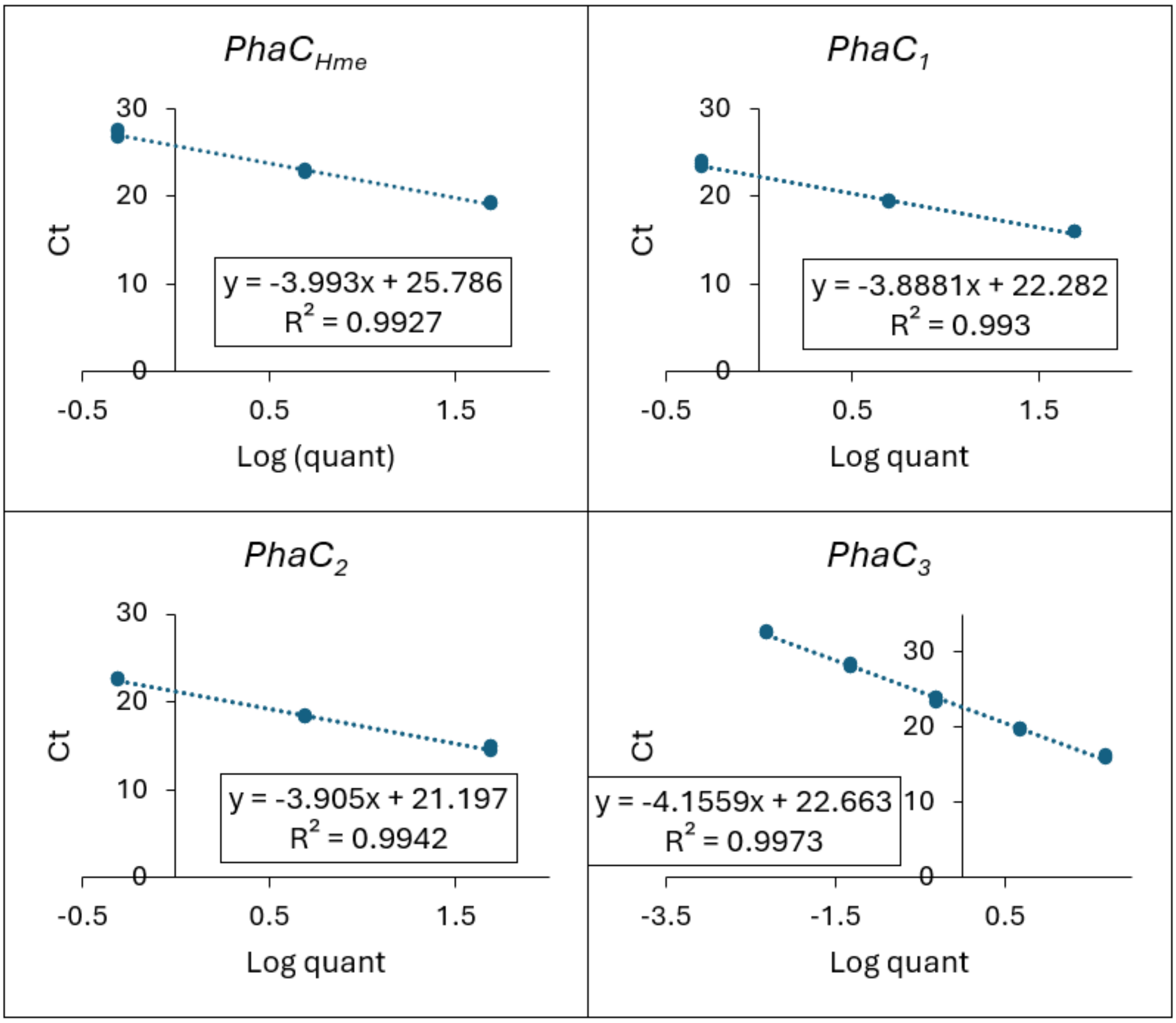


**Supplementary Figure S1:** Standard curves relating measured Ct values to input DNA quantities for all paralogs, measured on gDNA. The equations of the standard curves are used for estimation of the transcript levels.

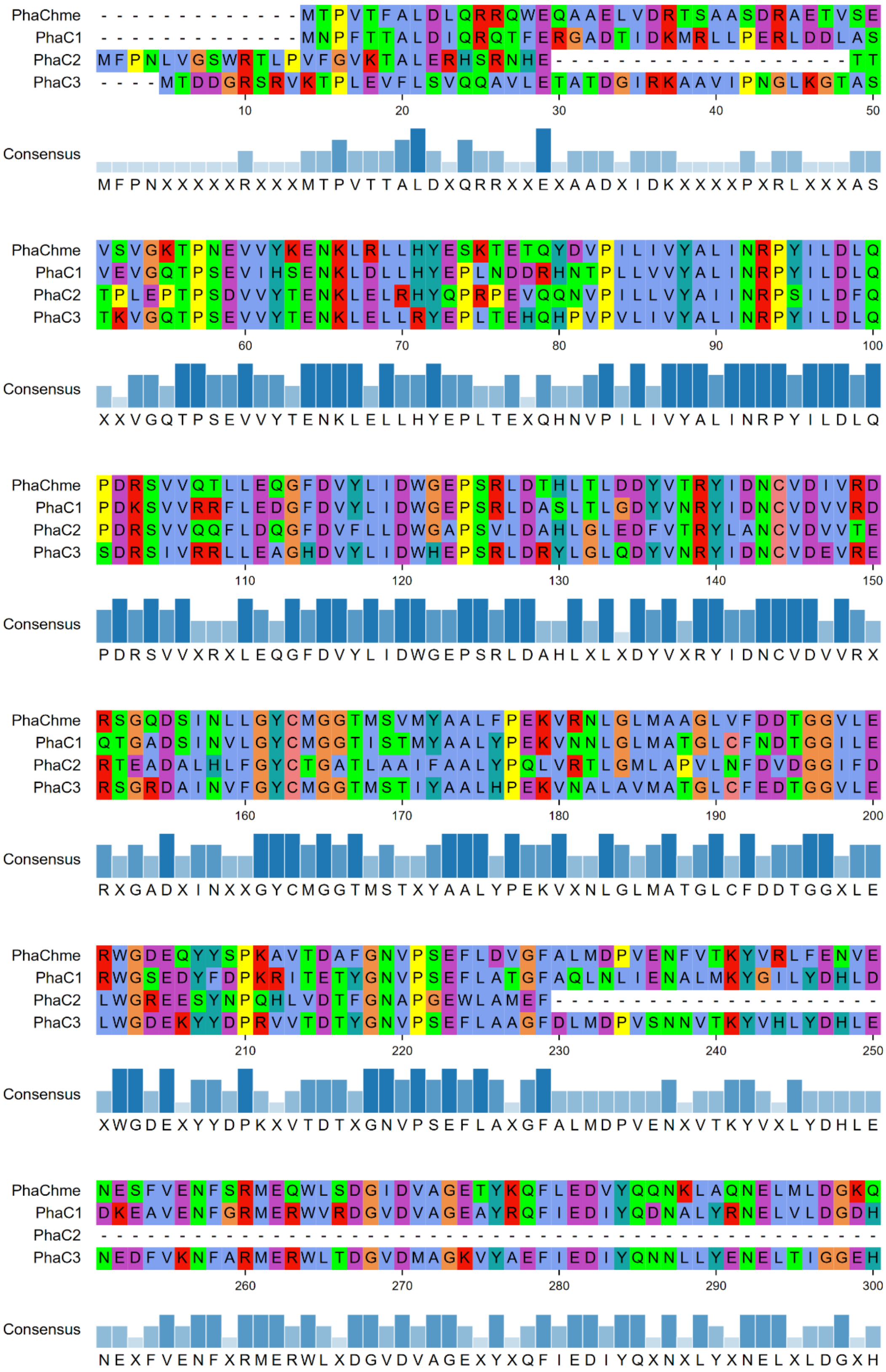

**Supplementary Figure S2:** Multiple sequence alignment of the four PhaC paralogs from *Haloferax mediterranei*. The alignment was visualized using pymsaviz with the Clustal color scheme. Consensus residues are indicated at the bottom of each panel. For clarity, the alignment is presented in two consecutive parts (1/2)

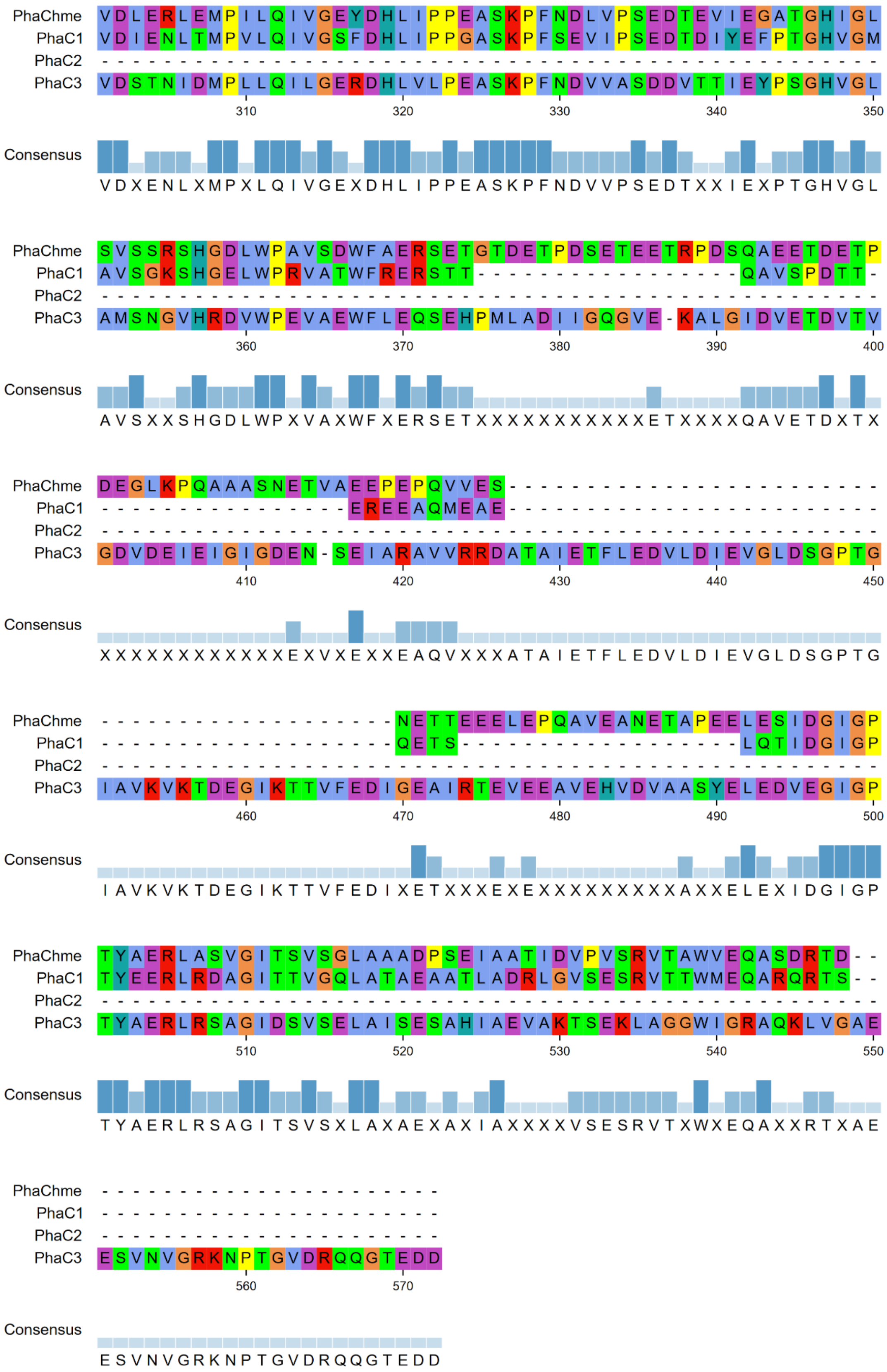


**Supplementary Figure S2:** Multiple sequence alignment of the four PhaC paralogs from *Haloferax mediterranei*. The alignment was visualized using pymsaviz with the Clustal color scheme. Consensus residues are indicated at the bottom of each panel. For clarity, the alignment is presented in two consecutive parts (2/2).

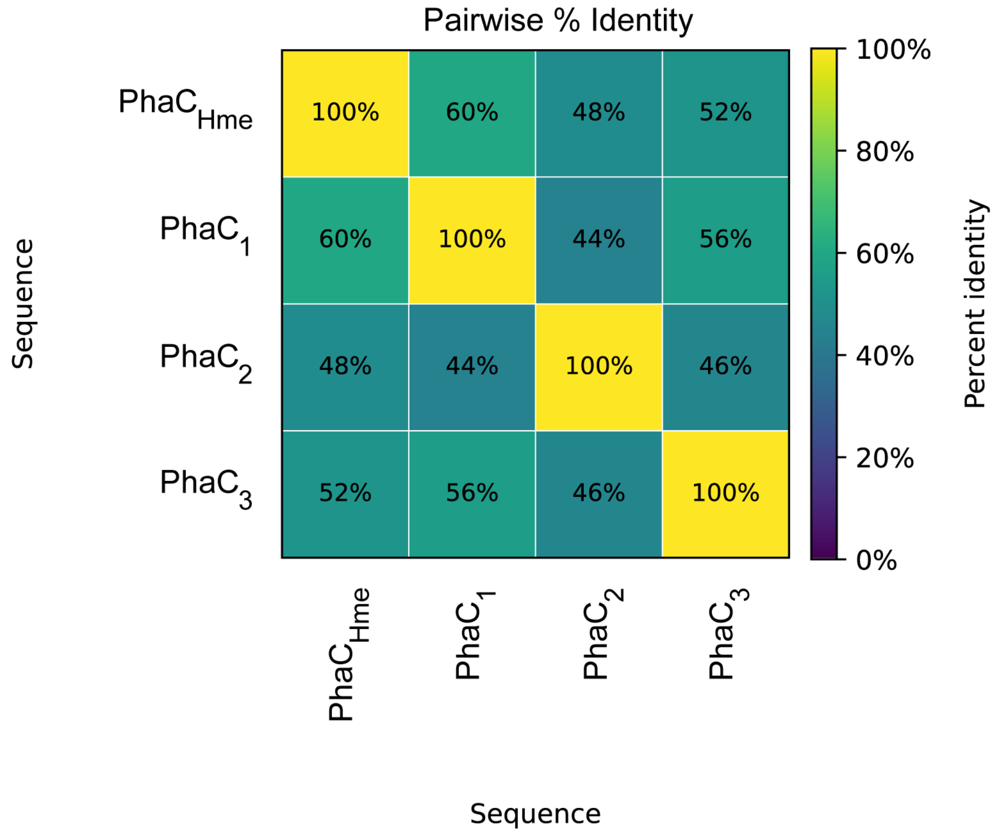


**Supplementary Figure S3:** Pairwise percentage identity matrix of aligned PhaC paralog sequences. Values represent the percentage of identical residues between each pair of sequences, calculated from the multiple sequence alignment with pairwise deletion of positions containing gaps.

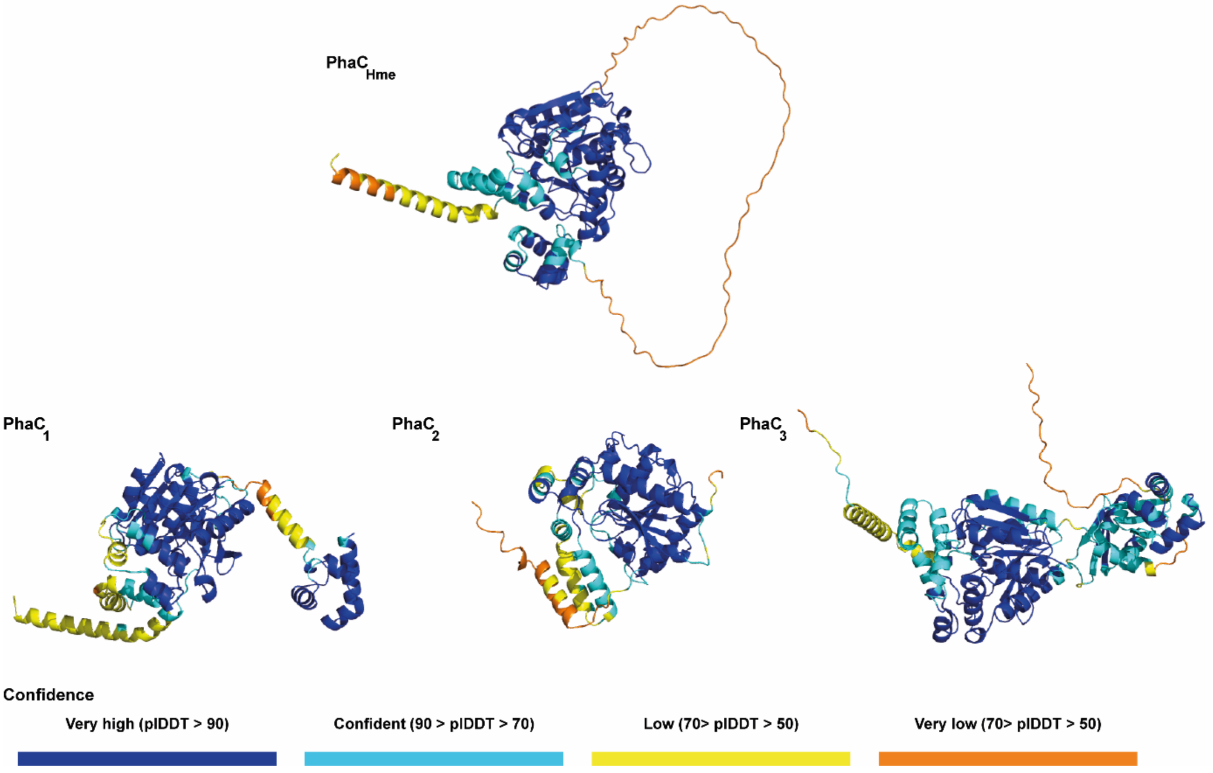


**Supplementary Figure S4:** AlphaFold models of $PhaC_{Hme}$ and its three paralogs. The structures are coloured based on their confidence scores according to the legend below.

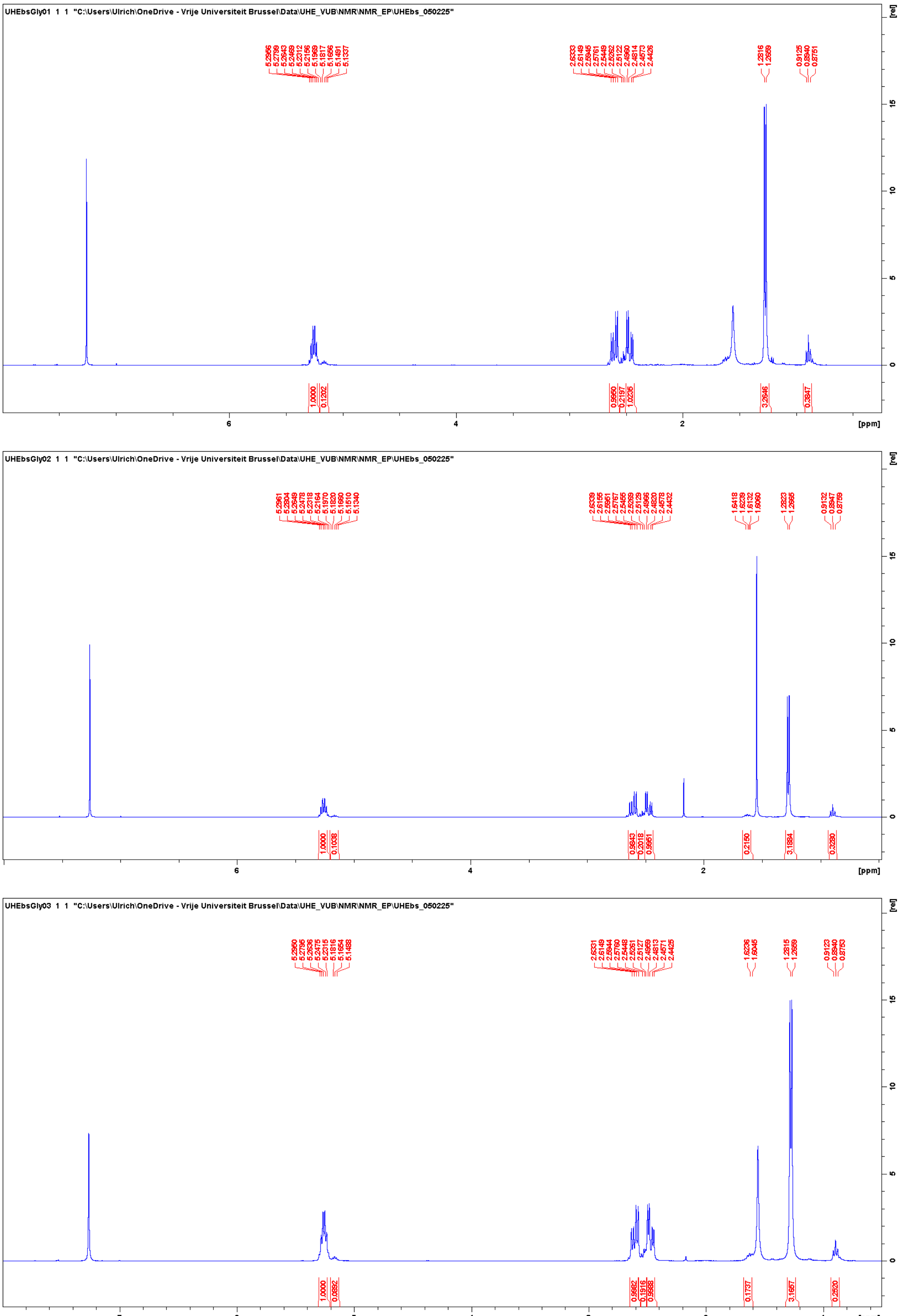


**Supplementary Figure S5:** $^{1}$H-NMR spectra of PHBV polymers extracted in triplicate from *Haloferax mediterranei* cells grown on the glycerol minimal medium.

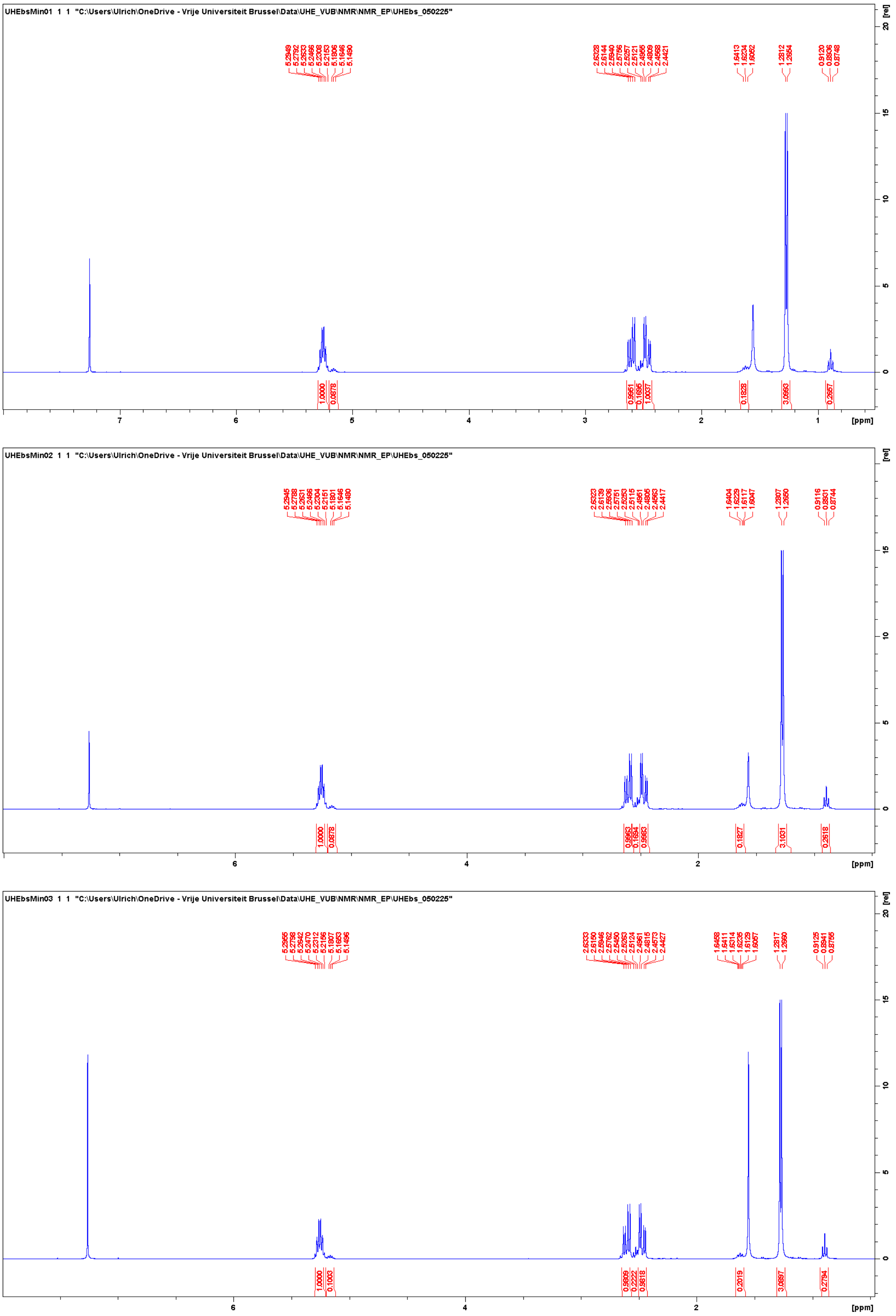

**Supplementary Figure S6:** ¹H-NMR spectra of PHBV polymers extracted in triplicate from *Haloferax mediterranei* cells grown on the Hv-min minimal medium.

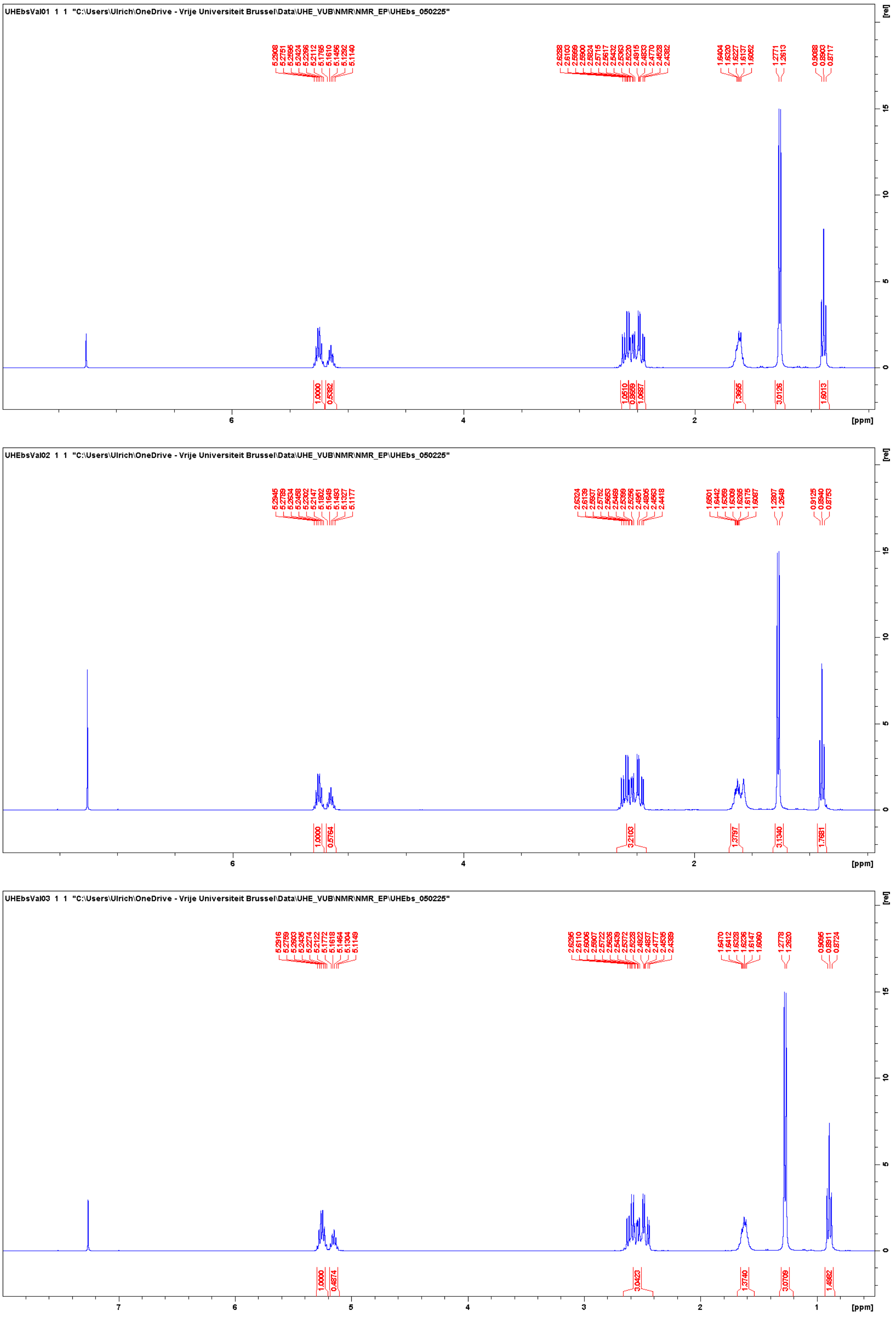


**Supplementary Figure S7:** $^{1}$H-NMR spectra of PHBV polymers extracted in triplicate from *Haloferax mediterranei* cells grown on the Hv-min minimal medium with the addition of valeric acid.

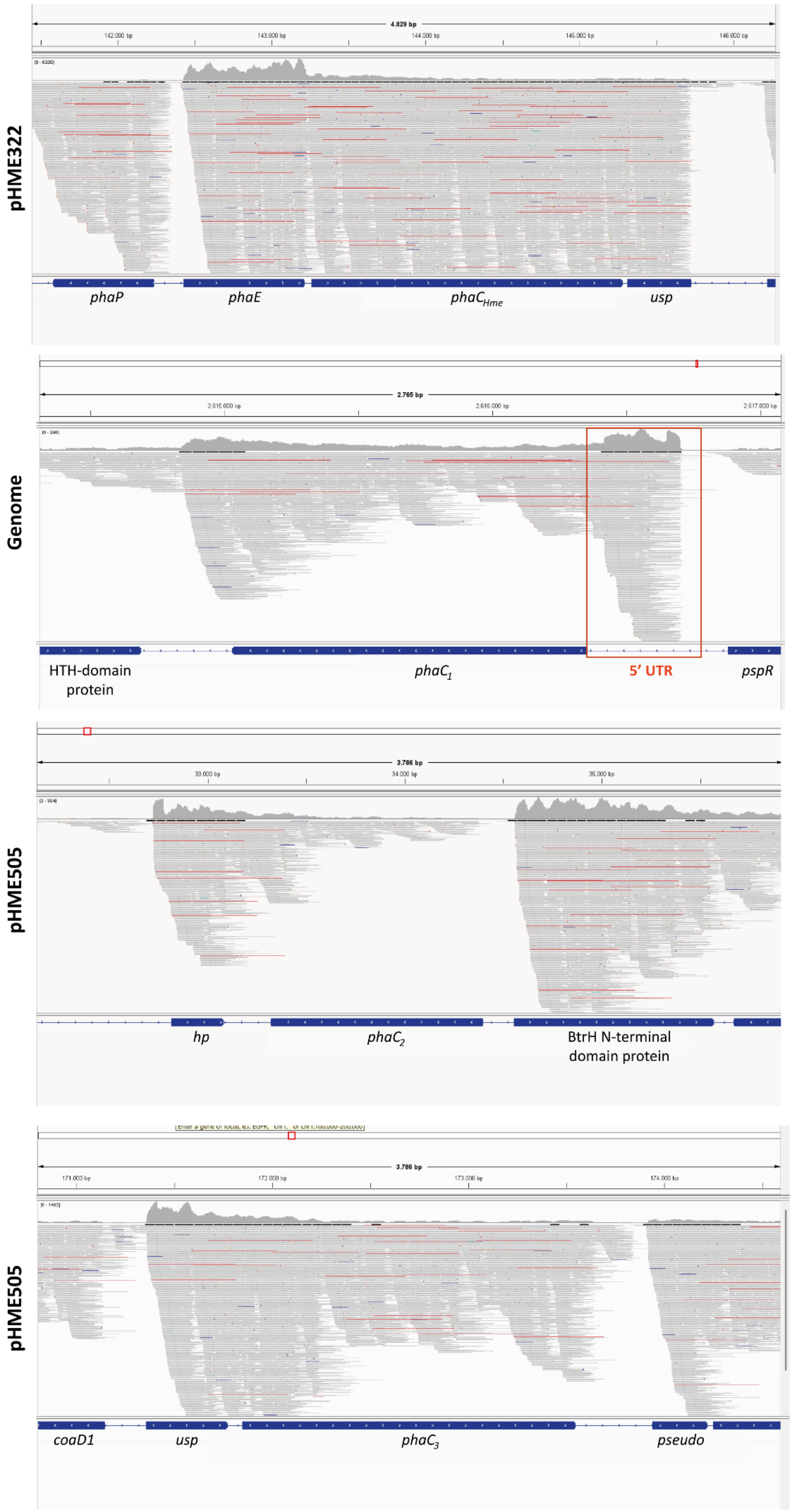


**Supplementary Figure S8:** Visualization of the mapped RNA-seq data published (Run: **SRR30916285**) previously by Martinez-Pastor *et al*. (2025) zoomed in on the genomic location of the paralogs. In red, the location of a hypothetical 5'UTR is boxed downstream of the *phaC$_1$* gene.

**Supplementary Table S1:** List of the genes monitored by RT-qPCR with their respective primers (forward F and reverse R) sequences and amplification efficiencies.

| Gene name (locus) | Gene abbreviation | F/R | Sequence (5' → 3') | Amplification efficiency (%) |
|---|---|---|---|---|
| TATA-box-binding protein (E6P09_03900) | *tbp* | F | ATCTCAACGCCATCGCAATCG | 80.2 |
| | | R | TACCCGACCCGAAGAGAAGTGC | |
| Signal recognition particle RNA (E6P09_08165) | *ffs* | F | AGTTAGGCCCTGCTCTTCACC | 68.1 |
| | | R | GGTTTCTACGTTGGCTTCCG | |
| PHA synthase subunit PhaC (E6P09_17995) | $phaC_{Hme}$ | F | GACTACGTGACTCGGTACATCG | 78.0 |
| | | R | CAGTACCCGAGAAGGTTAATCG | |
| PHA synthase subunit PhaC1 (E6P09_13380) | $phaC_1$ | F | CAACAATCTCGGACTCATGG | 80.8 |
| | | R | TCCGTACGTTTCGGTAATCC | |
| PHA synthase subunit PhaC2 (E6P09_15240) | $phaC_2$ | F | CCGCTATCTCAGCCTCTACG | 80.3 |
| | | R | TGTCAAAACCCCACTGAAGC | |
| PHA synthase subunit PhaC3 (E6P09_15825) | $phaC_3$ | F | CTCGCTATCTCCGAAAGTGC | 74.0 |
| | | R | TGGGGTTCTTTCTACCAACG | |

**Supplementary Table S2:** Overview of equations of standard curves of all paralogs, used for estimation of transcript levels.

| Gene abbreviation | Equation of standard curve |
|---|---|
| $phaC_{Hme}$ | Ct = -3.993 · log(quant) + 25.786 |
| $phaC_1$ | Ct = -3.888 · log(quant) + 22.282 |
| $phaC_2$ | Ct = -3.905 · log(quant) + 21.197 |
| $phaC_3$ | Ct = -4.156 · log(quant) + 22.663 |
